# Test-retest reliability of multi-metabolite edited MRS at 3T using PRESS and sLASER

**DOI:** 10.1101/2025.06.07.657685

**Authors:** Jessica Archibald, Amy E. Bouchard, Ralph Noeske, Dikoma C. Shungu, Mark Mikkelsen

## Abstract

**Purpose:** Spectral editing is the most common MRS approach for noninvasive in vivo measurement of low-concentration, strongly overlapped metabolites in the brain, such as γ-aminobutyric acid (GABA) and glutathione (GSH). Multi-metabolite editing methods, including HERMES and HERCULES, have recently been introduced, where multiple *J*-coupled metabolites can be edited in a single acquisition without increasing total scan time. Yet little is known regarding the reliability of these methods. This study assessed the test-retest reliability of HERMES and HERCULES, where volume localization was achieved using either PRESS or sLASER.

**Methods:** Sixteen healthy adult volunteers were scanned twice in two separate sessions. Single-voxel edited MRS data were acquired in the medial parietal lobe using the following sequences: (1) HERMES-PRESS; (2) HERMES-sLASER; (3) HERCULES-PRESS; (4) HERCULES-sLASER. Spectra were processed and metabolites were quantified using the Osprey software. Data quality metrics and reliability statistics were estimated for all four acquisitions.

**Results:** HERMES-sLASER demonstrated lower within-subjects coefficients of variation (CV_ws_) for GSH, glutamine (Gln), and glutamate (Glu) + Gln (Glx), suggesting improved reliability compared to HERMES-PRESS. However, GABA + co-edited macromolecules (GABA+) and Glu showed higher CV_ws_ for HERMES-sLASER. HERCULES-sLASER produced better reliability than HERCULES-PRESS for GABA+, GSH, Glu, Gln, Glx, aspartate (Asp), and lactate (Lac). *N*-acetylaspartate (NAA) and *N*-acetylaspartylglutamate (NAAG) showed higher CV_ws_ for HERCULES-sLASER. These findings suggest that sLASER may be more advantageous than PRESS for volume localization in simultaneous multi-metabolite editing.

**Conclusion:** Using sLASER yielded better test-retest reliability for most metabolites than using PRESS for volume localization for HERMES and HERCULES.

## Introduction

Magnetic resonance spectroscopy (MRS) is a noninvasive biomedical spectroscopic technique that enables the measurement and quantification of a wide range of metabolites in vivo. To better understand the brain, research has investigated the involvement of these metabolites in sensory stimulation^1–5^, several neurological and psychiatric disorders^6–11^, and healthy aging^12^. However, multiple metabolite resonances, some with similar chemical shifts, result in substantial spectral overlap, making peak assignment and quantification of detected metabolites in vivo challenging. The set of methods designed to address this is collectively referred to as spectral editing^13^. Two highly investigated editable metabolites are γ-aminobutyric acid (GABA) and glutathione (GSH) due to their relevance in aging^14,15^, cancer^16,17^, cognition^18^, psychiatric disorders^19–21^, motor performance^18^, and neurological disorders^22,23^. However, a disadvantage is that only a single metabolite is typically targeted in each acquisition. In addition to the relatively long scan times required to improve SNR, single-metabolite editing can be inefficient because other metabolites of interest may not be measured within a scan session. Recently, novel editing methods, namely Hadamard Encoding and Reconstruction of MEGA-Edited Spectroscopy (HERMES)^24–26^ and Hadamard Editing Resolves Chemicals Using Linear-Combination Estimation of Spectra (HERCULES)^27^, were introduced where multiple *J*-coupled metabolites can be targeted in a single acquisition without increasing total scan time. HERMES has mostly been used to target GABA and GSH, while HERCULES is designed to examine several more metabolites; that is, GABA, GSH, ascorbate (Asc), aspartate (Asp), NAA, *N-*acetylaspartylglutamate (NAAG) and lactate (Lac) (2-hydroxyglutarate (2HG) is also targetable in brain tumors). This is achieved through multiplexed editing schemes (with four or more shots) and Hadamard reconstruction. However, the reliability of these multi-metabolite editing techniques has yet to be investigated, especially in combination with different localization techniques.

Restricting the signal detection to a well-defined region of interest is crucial for all MRS techniques. The goal is to remove all unwanted signals outside the region, such as lipids, and manage tissue (i.e., gray matter, white matter, and CSF) differences, as each tissue has its unique metabolic profile^13^. PRESS^28^ localization is most commonly used and has been the basis for introducing novel multi-metabolite editing techniques^24,26,27^. However, using PRESS-based slice-selective RF pulses introduces chemical shift displacement errors (CSDEs) due to differences in chemical shifts between resonances and the bandwidths of the pulses. These CSDEs can lead to significant discrepancies in the localization of various resonances. In contrast, semi-localized by adiabatic selective refocusing (sLASER)^29^ is a sequence increasingly recognized as a theoretically more reliable localization method^30^. The adiabatic RF pulses that sLASER uses allow localization to be better defined, resulting in lower CSDEs. sLASER is also less sensitive to *B*_1_ inhomogeneity than PRESS^31^.

This study compared the test-retest reliability of HERMES and HERCULES implemented with sLASER and PRESS for volume localization. Due to the benefits of sLASER, it was hypothesized that sLASER would improve the reliability of the detection and quantification of various metabolites.

## Methods

### Participants

Sixteen healthy adult volunteers (male/female = 6/10; mean age ± 1 std = 38.4 ± 18.2 years) were recruited for this study. Each volunteer underwent two scan sessions separated by a time delay, with a median interval between scans of 0 days (range: 0–29 days). Participants were excluded if they had any contraindications for MRI or a history of neurological or psychiatric disorders. The Weill Cornell Medicine Institutional Review Board granted ethical approval for this study. All participants provided written informed consent before taking part in the study.

### MR scanning protocol

All data were collected on a 3T GE Discovery MR750 MRI scanner using a ^1^H 32-channel RF phased-array head coil for receive and a body coil for transmit.

#### MRI

High-resolution 3D *T*_1_-weighted BRAVO structural scans (FSPGR; TE/TR/TI = 5.2/12.2/725 ms; flip angle = 7°; voxel resolution = 0.9 × 0.9 × 1.5 mm^3^; matrix size = 256 × 256; slices = 124; parallel acceleration factor = 2) were first acquired for accurate voxel placement in each scan session.

#### MRS

Single-voxel multi-metabolite-edited MRS data were acquired in the following order (and were not counterbalanced between sessions): (1) HERMES-PRESS; (2) HERMES-sLASER; (3) HERCULES-PRESS; (4) HERCULES-sLASER; with the following parameters TE/TR = 82/2000 ms; spectral width = 5000 Hz; 4096 points; 224 transients; and voxel resolution = 3 × 3 × 3 cm^3^. The MRS voxel was placed in the medial parietal lobe (**Figure 1**). PRESS-localized scans used CHESS for water suppression, while sLASER scans used VAPOR. Although this is a methodological discrepancy, VAPOR is currently the recommended water suppression method^32^, while GE has used CHESS for PRESS by default for decades. Therefore, we decided to use the standards that matched PRESS and sLASER acquisitions to align with ecological validity (i.e., what most users would most usually implement). An MRSinMRS checklist^33^ for this study is provided in **Table S1**.

**Figure 1.**
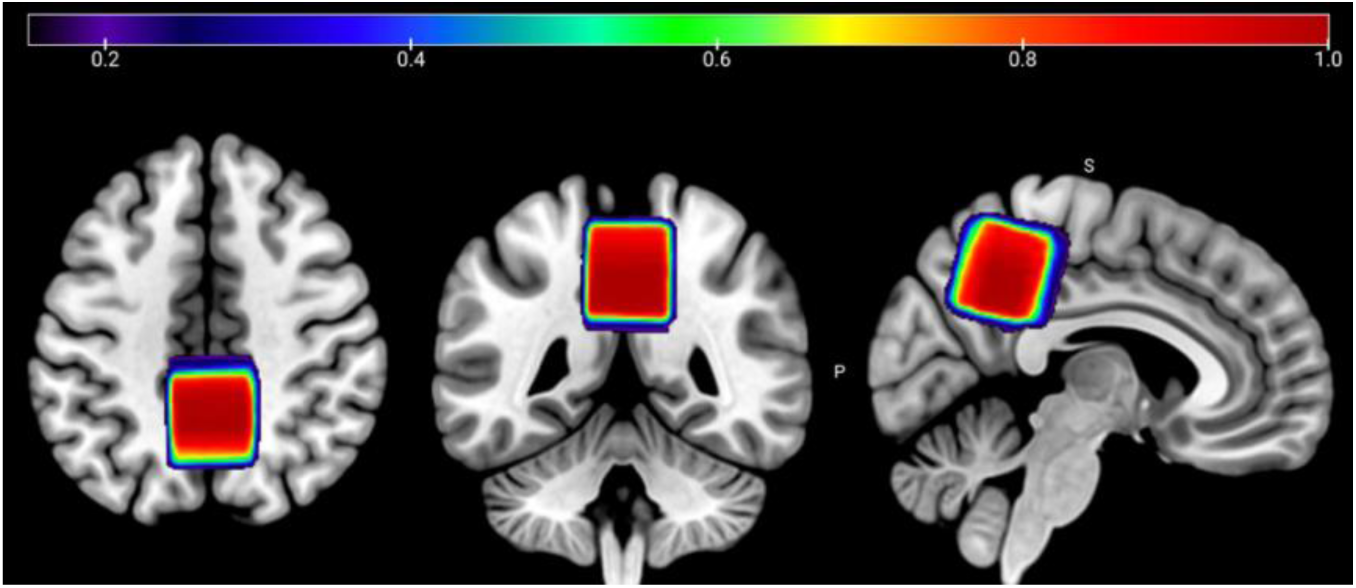
Voxel placement map showing the average overlap between scan sessions 1 and 2 in the medial parietal lobe for all participants in MNI152 template space. The color bar denotes the estimated overlap, where 1.0 equates to 100% overlap.

### Data analysis

Spectra were processed using Osprey^34^ (v2.5.0) and involved the following steps: (1) RF coil combination using generalized least squares^35^; (2) eddy-current correction^36^; (3) robust spectral registration^37^; (4) signal averaging; (5) residual water filtering using Hankel singular value decomposition^38^; and (6) reconstruction of four Hadamard combinations based on the four subspectra labeled A, B, C, and D^25,27^. The resulting subspectra were combined to make three combinations of interest: (1) the SUM spectrum (A+B+C+D); (2) the DIFF1 spectrum (A+B–C– D); and (3) the DIFF2 spectrum (A–B+C–D).

Reconstructed spectra were fitted using nonlinear least-squares linear-combination modeling. Basis sets were created using high spatial resolution (101 × 101 points) density-matrix numerical simulations. These were run using the HERMES- and HERCULES-edited PRESS and sLASER pulse sequence parameters to accurately simulate metabolite signal line shapes. Simulations were run in a customized version of MRSCloud^39^ using a 1D projection method^40^ and coherence pathway filtering^41^ to reduce computation time. Metabolites included in the basis sets for HERMES and HERCULES were Asc, Asp, Cr, negative creatine methylene (-CrCH_2_), GABA, Gln, Glu, glycerophosphocholine, GSH, H_2_O, Lac, NAA, NAAG, *myo*-inositol (mI), phosphorylcholine, PCr, phosphoethanolamine, *scyllo*-inositol, and taurine. Macromolecule and lipid resonances were parameterized using Gaussian functions.

SNR was calculated as the ratio between the fitted 3 ppm Cr peak model amplitude in the SUM spectrum and the standard deviation of the noise signal between –2 and 0 ppm. Linewidth was determined by the FWHM of the model fit of the unsuppressed water signal in the frequency domain. The model fit errors of the SUM, DIFF1, and DIFF2 spectra were estimated as the sum of squares of residuals normalized to the square of the standard deviation of the noise signal between –2 and 0 ppm and multiplied by the number of points of the residuals.

### Quantification

The metabolites of interest for HERMES and HERCULES were quantified using unsuppressed water as a reference signal and are reported in institutional units (i.u.). No corrections for partial-volume tissue effects were applied. For the HERMES acquisitions, GABA+, Glu, Gln, and Glx were quantified from the DIFF1 spectrum, while GSH was quantified from the DIFF2 spectrum. For the HERCULES acquisitions, GABA+, Glu, Gln, and Glx were quantified from the DIFF1 spectrum, GSH, Asp, Lac, and NAAG were quantified from the DIFF2 spectrum, and NAA was quantified from the SUM spectrum. Asc was not detectable during fitting and was consequentially excluded from further analysis.

### Statistical analysis

All statistical analyses were performed in R (v4.4.0). Multivariate outliers were removed using the robust Mahalanobis-minimum covariance determinant (MMCD) distance with a quantile of 0.75 and an alpha of 0.01^42^. Coefficients of variation (CV) were calculated for both within-subjects (CV_ws_) and between-subjects (CV_bs_). CV_ws_ were calculated using the root-mean-squared approach. Statistical differences in spectral data quality metrics were assessed using two-way repeated-measures ANOVA. Post hoc comparisons were conducted using Tukey’s honest significant difference method with *p*-values adjusted to account for multiple comparisons.

## Results

One volunteer had no HERMES data for either session, and another had no usable HERCULES-sLASER data for either session due to incorrect acquisition parameters. Additionally, two data sets for HERCULES-sLASER and HERCULES-PRESS for session 2 were not obtained, as the respective volunteers opted to discontinue the session before its completion. All spectra underwent visual inspection for data quality and signal artifacts. Subsequently, HERCULES-PRESS data from one volunteer from session 2 was excluded due to an unstable baseline, and one participant’s HERMES-PRESS data from session 1 was excluded due to poor frequency-and-phase alignment of transients.

Figures 2 and **3** show sample data from one participant for all acquisition schemes and reconstructed subspectra (SUM, DIFF1, and DIFF2), including basis set functions for quantified metabolites.

**Figure 2.**
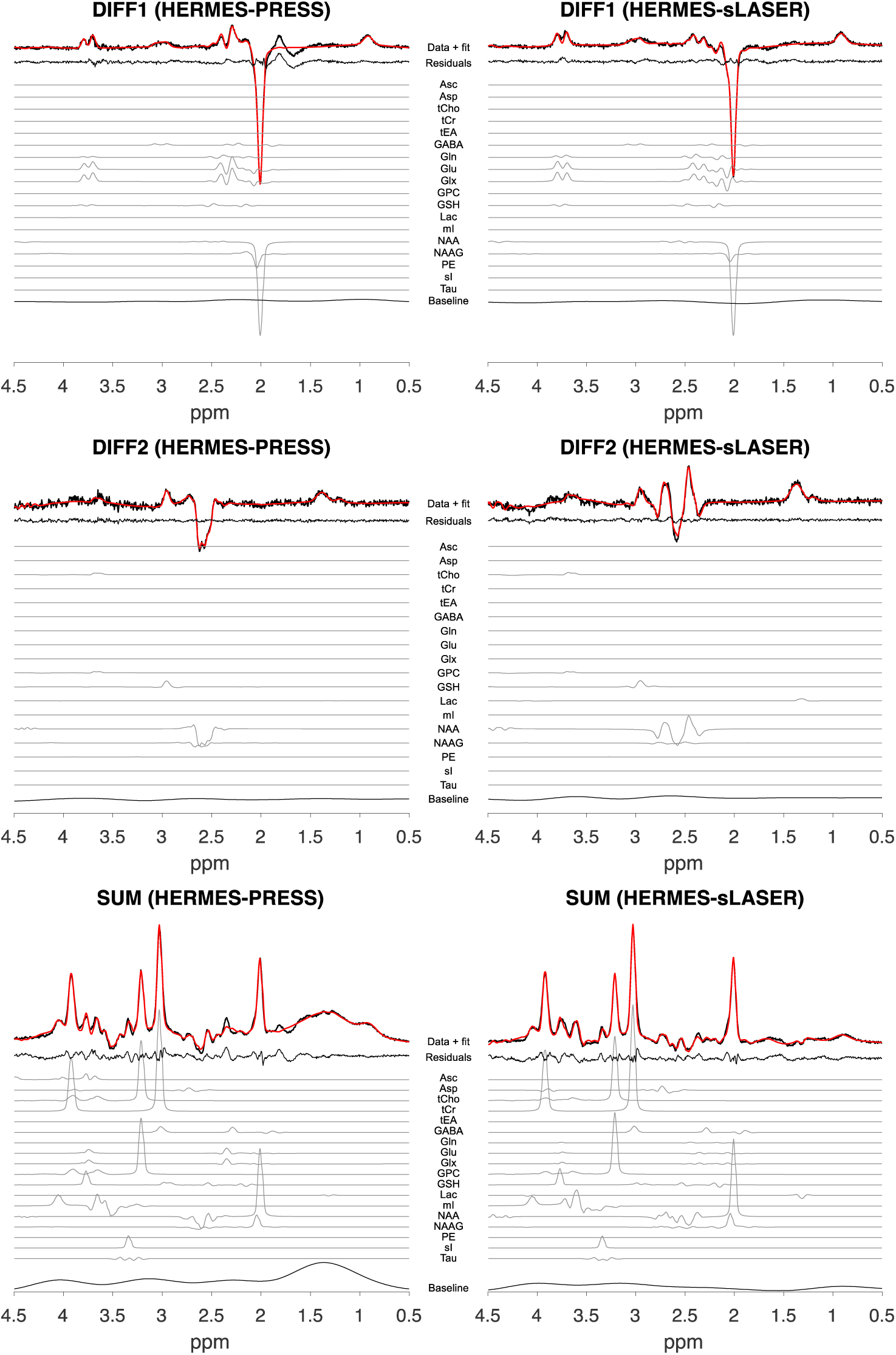
Example HERMES-PRESS and HERMES-sLASER spectra from one participant. Spectral data for each Hadamard combination (DIFF1, DIFF2, and SUM) (in black) and corresponding model fits (in red) are shown. The model fit residuals are also plotted. Additionally, the individual metabolite basis set fits are displayed (not to scale), with the corresponding metabolites of interest highlighted in color. At the bottom of each subplot is the baseline signal fit.

**Figure 3.**
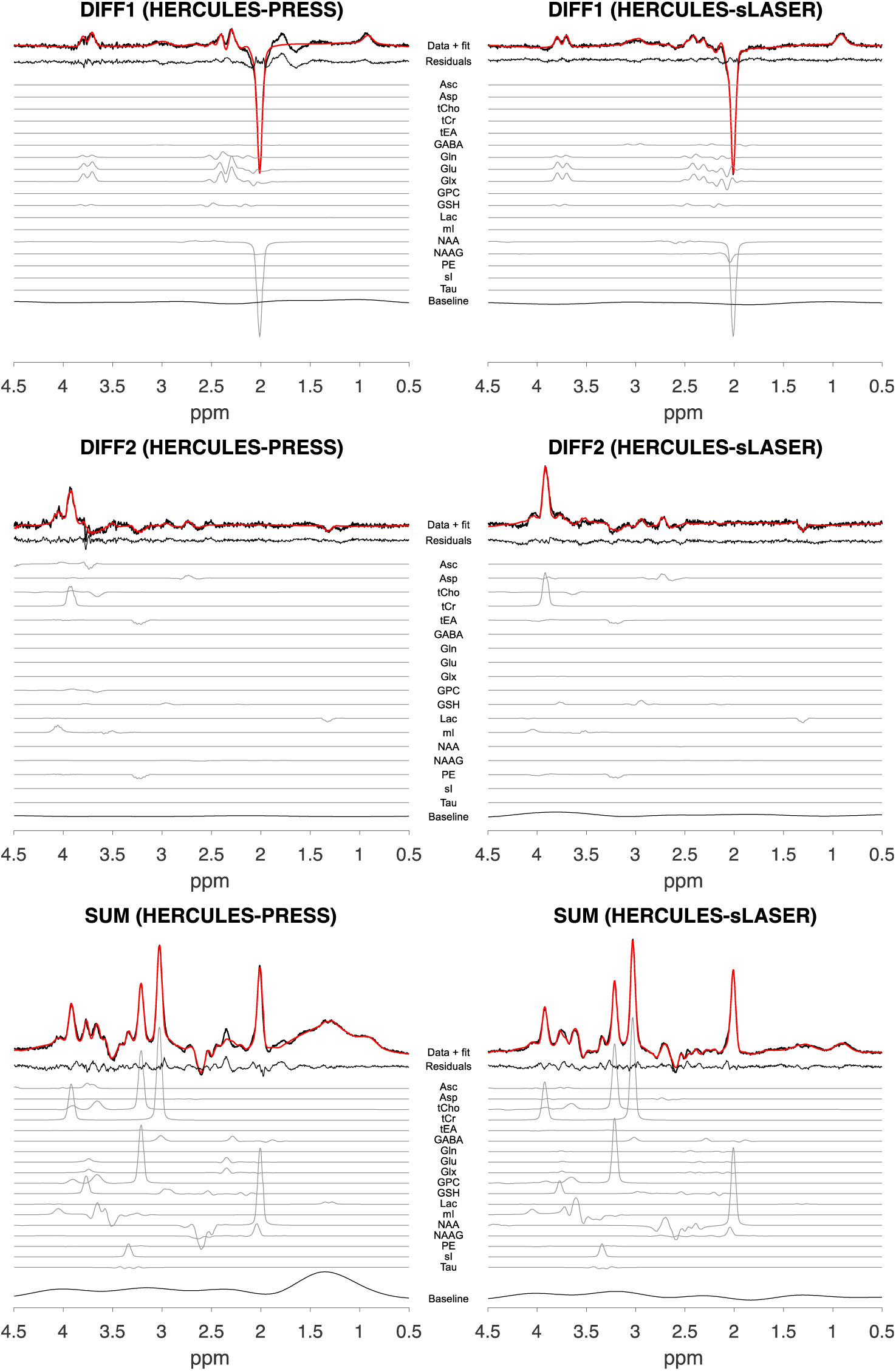
Example HERCULES-PRESS and HERCULES-sLASER spectra from the same participant as in Figure 2.

### Spectral data quality metrics

For the remaining non-excluded participants, **Table 1** details the average Cr SNR, unsuppressed water linewidth (FWHM), and fit error of each reconstructed spectrum. **Table 2** shows the results of the analyses with respect to statistical differences in the data quality metrics. There was no significant difference in SNR across scan sessions (*p* = 0.39) or acquisitions (*p* = 0.70). There was no significant difference in linewidth across sessions (*p* = 0.09). On the other hand, there was a significant linewidth difference across acquisitions (*p =* 0.013). Post hoc comparisons revealed that HERMES-sLASER had a lower linewidth than HERMES-PRESS (*p =* 0.025). HERCULES-sLASER also had a lower linewidth than HERCULES-PRESS (*p* = 0.029). In addition, HERMES-PRESS had a larger linewidth than HERCULES-sLASER (*p =* 0.03). Lastly, the linewidth was lower for HERMES-sLASER compared to HERCULES-PRESS (*p =* 0.025).

**Table 1.**
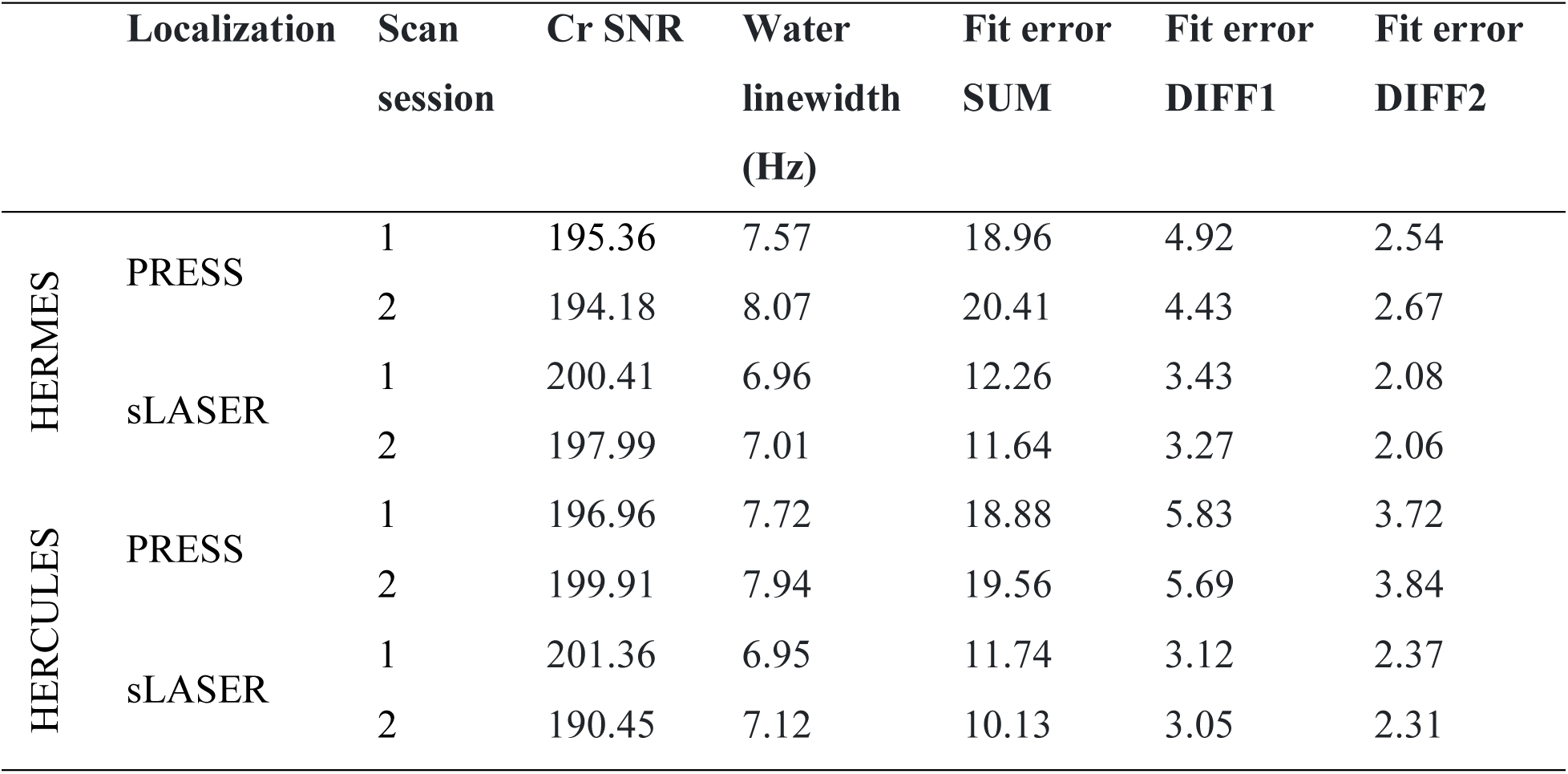
Mean spectral data quality metrics.

|  | Localization | Scan session | Cr SNR | Water linewidth (Hz) | Fit error SUM | Fit error DIFF1 | Fit error DIFF2 |
| --- | --- | --- | --- | --- | --- | --- | --- |
| HERMES | PRESS | 1 | 195.36 | 7.57 | 18.96 | 4.92 | 2.54 |
|  |  | 2 | 194.18 | 8.07 | 20.41 | 4.43 | 2.67 |
|  | sLASER | 1 | 200.41 | 6.96 | 12.26 | 3.43 | 2.08 |
|  |  | 2 | 197.99 | 7.01 | 11.64 | 3.27 | 2.06 |
| HERCULES | PRESS | 1 | 196.96 | 7.72 | 18.88 | 5.83 | 3.72 |
|  |  | 2 | 199.91 | 7.94 | 19.56 | 5.69 | 3.84 |
|  | sLASER | 1 | 201.36 | 6.95 | 11.74 | 3.12 | 2.37 |
|  |  | 2 | 190.45 | 7.12 | 10.13 | 3.05 | 2.31 |

**Table 2.** ANOVA results and post hoc comparisons for the data quality metrics.

| **SNR** |  |  |  |
| --- | --- | --- | --- |
| ***ANOVA*** |  |  |  |
|  | *df* | *F*-value | *p* |
| Session | 1 | 0.99 | 0.39 |
| Acquisition | 3 | 0.51 | 0.70 |
| Residuals | 3 |  |  |
| **Linewidth** |  |  |  |
| ***ANOVA*** |  |  |  |
|  | *df* | *F*-value | *p* |
| Session | 1 | 6.09 | 0.09 |
| Acquisition | 3 | 24.45 | **0.013** |
| Residuals | 3 |  |  |
| ***Post hoc comparisons*** |  |  |  |
|  |  | Difference | *p* (adj)<sup>a</sup> |
| Session 2 – Session 1 |  | 0.24 | 0.09 |
| HERCULES-sLASER – HERCULES-PRESS |  | –0.80 | **0.029** |
| HERMES-PRESS – HERCULES-PRESS |  | –0.01 | > 0.99 |
| HERMES-sLASER – HERCULES-PRESS |  | –0.85 | **0.025** |
| HERMES-PRESS – HERCULES-sLASER |  | 0.79 | **0.03** |
| HERMES-sLASER – HERCULES-sLASER |  | –0.05 | 0.98 |
| HERMES-sLASER – HERMES-PRESS |  | –0.84 | **0.025** |
| **Fit error: SUM** |  |  |  |
| ***ANOVA*** |  |  |  |
|  | *df* | *F*-value | *p* |
| Session | 1 | 0.001 | 0.97 |
| Acquisition | 3 | 43.19 | **0.005** |
| Residuals | 3 |  |  |

*Post hoc comparisons*
|  | Difference | *p* (adj) |
| --- | --- | --- |
| Session 2 – Session 1 | –0.025 | 0.97 |
| HERCULES-sLASER – HERCULES-PRESS | –8.29 | **0.01** |
| HERMES-PRESS – HERCULES-PRESS | 0.47 | 0.96 |
| HERMES-sLASER – HERCULES-PRESS | –7.27 | **0.014** |
| HERMES-PRESS – HERCULES-sLASER | 8.75 | **0.009** |
| HERMES-sLASER – HERCULES-sLASER | 1.015 | 0.74 |
| HERMES-sLASER – HERMES-PRESS | –7.74 | **0.012** |

**Fit error: DIFF1***ANOVA*
|  | *df* | *F*-value | *p* |
| --- | --- | --- | --- |
| Session | 1 | 5.27 | 0.11 |
| Acquisition | 3 | 175.64 | < **0.001** |
| Residuals | 3 |  |  |

*Post hoc comparisons*
|  | Difference | *p* (adj) |
| --- | --- | --- |
| Session 2 – Session 1 | –0.22 | 0.11 |
| HERCULES-sLASER – HERCULES-PRESS | –2.68 | < **0.001** |
| HERMES-PRESS – HERCULES-PRESS | –1.09 | **0.012** |
| HERMES-sLASER – HERCULES-PRESS | –2.41 | **0.001** |
| HERMES-PRESS – HERCULES-sLASER | 1.59 | **0.004** |
| HERMES-sLASER – HERCULES-sLASER | 0.27 | 0.35 |
| HERMES-sLASER – HERMES-PRESS | –1.33 | **0.007** |

**Fit error: DIFF2***ANOVA*
|  | *df* | *F*-value | *p* |
| --- | --- | --- | --- |
| Session | 1 | 0.77 | 0.44 |
| Acquisition | 3 | 242.48 | < **0.001** |

Residuals 3 ***Post hoc comparisons***
|  | Difference | *p* (adj) |
| --- | --- | --- |
| Session 2 – Session 1 | 0.043 | 0.44 |
| HERCULES-sLASER – HERCULES-PRESS | –1.44 | **< 0.001** |
| HERMES-PRESS – HERCULES-PRESS | –1.18 | **0.001** |
| HERMES-sLASER – HERCULES-PRESS | –1.71 | **< 0.001** |
| HERMES-PRESS – HERCULES-sLASER | 0.27 | 0.088 |
| HERMES-sLASER – HERCULES-sLASER | –0.27 | 0.084 |
| HERMES-sLASER – HERMES-PRESS | –0.54 | **0.013** |

<sup>a</sup> *p*-value adjusted for multiple comparisons based on Tukey’s honest significant difference method.

Across sessions, there were no significant differences in fit error for SUM (*p =* 0.97), DIFF1 (*p =* 0.11), or DIFF2 (*p =* 0.44). However, there were significant differences across acquisitions for all three: SUM (*p =* 0.005), DIFF1 (*p* < 0.001), and DIFF2 (*p* < 0.001). Post hoc comparisons of fit error for the three reconstructed spectra are displayed in **Table 2**.

### Test-retest reliability

**Table 3** summarizes the test-retest reliability results for each editing technique and localization method. The mean water-referenced concentration estimates for each metabolite of interest and their respective CV_ws_ and CV_bs_ are reported. Figure 4 depicts Bland-Altman plots displaying the agreement between scan sessions for GABA+, GSH for HERMES and GABA+, GSH, Asp, Lac, NAA, and NAAG for HERCULES. These plots are presented as the average of the measurements across scan sessions versus the average difference between scan sessions as a percentage. This selection of metabolites of interest reflects the targets of the GABA-GSH HERMES^25^ and HERCULES publications^27^.

**Figure 4.**
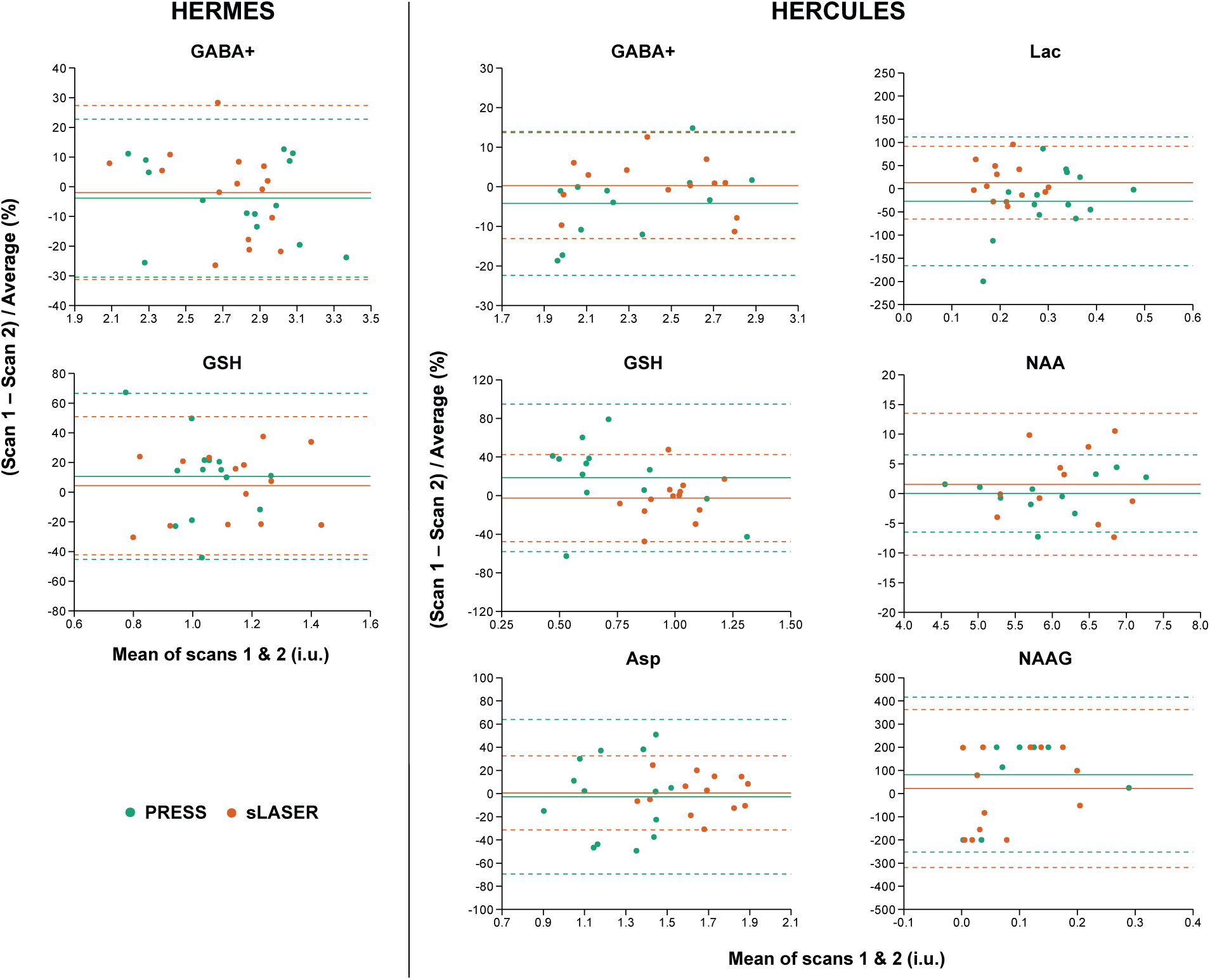
Bland-Altman plots for the metabolites of interest. GABA+ and GSH are displayed for HERMES on the left side of the figure. GABA+, GSH, Asp, Lac, NAA, and NAAG are shown for HERCULES towards the right. PRESS and sLASER data are shown for each metabolite in green and orange, respectively.

**Table 3.** Test-retest reliability statistics of multi-metabolite-edited PRESS and sLASER metabolite concentration estimates over scan sessions 1 and 2. The text in bold indicates better reliability between localization approaches within editing approach.

|  |  | Localization | Mean<br>concentration (i.u.) | CV<sub>ws</sub> | CV<sub>bs</sub> |
| --- | --- | --- | --- | --- | --- |
| **GABA+** | HERMES | PRESS | 2.78 | **9.60%** | 15.20% |
|  |  | sLASER | 2.77 | 10.60% | **10.30%** |
|  | HERCULES | PRESS | 2.30 | 7.00% | 14.30% |
|  |  | sLASER | 2.43 | **4.60%** | **13.30%** |
| **GSH** | HERMES | PRESS | 1.06 | 17.20% | **14.90%** |
|  |  | sLASER | 1.10 | **16.50%** | 20.10% |
|  | HERCULES | PRESS | 0.68 | 29.40% | 34.00% |
|  |  | sLASER | 0.99 | **15.70%** | **16.20%** |
| **Glu** | HERMES | PRESS | 6.58 | **5.40%** | 10.10% |
|  |  | sLASER | 6.97 | 7.30% | **8.40%** |
|  | HERCULES | PRESS | 6.22 | 10.00% | 15.50% |
|  |  | sLASER | 5.82 | **7.30%** | **12.40%** |
| **Gln** | HERMES | PRESS | 1.33 | 30.40% | 28.20% |
|  |  | sLASER | 1.61 | **18.60%** | **20.20%** |
|  | HERCULES | PRESS | 1.89 | 19.00% | 26.60% |
|  |  | sLASER | 1.79 | **14.60%** | **15.30%** |
| **Glx** | HERMES | PRESS | 7.93 | 7.70% | 10.60% |
|  |  | sLASER | 8.61 | **7.20%** | **8.00%** |
|  | HERCULES | PRESS | 8.16 | 13.30% | 18.50% |
|  |  | sLASER | 7.71 | **10.10%** | **13.60%** |
| **Asp** | HERCULES | PRESS | 1.26 | 23.30% | 22.90% |
|  |  | sLASER | 1.66 | **11.10%** | **13.20%** |
| **Lac** | HERCULES | PRESS | 0.31 | 52.00% | 38.50% |
|  |  | sLASER | 0.21 | **28.80%** | **30.10%** |
| **NAA** | HERCULES | PRESS | 5.93 | **2.20%** | 13.40% |
|  |  | sLASER | 6.26 | 5.70% | **10.20%** |
| **NAAG** | HERCULES | PRESS | 0.18 | **113.40%** | **100.40%** |
|  |  | sLASER | 0.06 | 125.80% | 173.70% |

i.u., institutional units; $CV_{ws}$ , within-subjects coefficient of variation; $CV_{bs}$ , between-subjects CV

HERMES-PRESS and HERMES-sLASER showed similar reliability for GABA+ and GSH. In comparison, HERCULES-sLASER had better GABA+ and GSH as well as Asp and Lac reliability than HERCULES-PRESS. NAA was more reliable using HERCULES-PRESS localization, while NAAG showed poor reliability for HERCULES-PRESS and HERCULES-sLASER.

Gln and Glx were more reliably measured with HERMES-sLASER and HERCULES-sLASER, while the reliability of Glu was better using HERMES-PRESS and HERCULES-sLASER.

## Discussion and Conclusions

This study aimed to evaluate the test-retest reliability of single-voxel multi-metabolite spectral-edited MRS at 3T using HERMES and HERCULES, in which volume localization was achieved with PRESS or sLASER. We hypothesized that sLASER, with its recognized benefits, would show improved reliability for detecting the targeted metabolites. Our study produced results suggesting an overall advantage of sLASER over PRESS localization for most metabolites of interest. HERMES-sLASER demonstrated lower CV_ws_ for GSH, Gln, and Glx and improved reliability compared to HERMES-PRESS. GABA+ and Glu, however, showed higher CV_ws_ for sLASER. HERCULES-sLASER had better reliability than HERCULES-PRESS for multiple metabolites, including GABA+, GSH, Glu, Gln, Glx, Asp, and Lac, while NAA and NAAG showed higher CV_ws_. However, NAAG for both HERCULES-PRESS and HERCULES-sLASER had very high CV_ws_ values.

Although not much is known regarding the reliability of sLASER (unlike PRESS^43–49^ and MEGA-PRESS ^44,49–54^), there is some evidence that sLASER is a better localization method. For instance, in one study, sLASER had better reliability than PRESS across multiple metabolites, such as mI, NAA + NAAG, *scyllo*-inositol, tCho, and tCr but not Glu, possibly from PRESS overestimating Glu concentrations^55^. Additionally, sLASER displayed improved reliability (higher ICCs) than STEAM for Glu at 7T^56^. An additional 7T study also showed that sLASER had better reliability (lower CV) than STEAM for mI but higher CV for GABA, potentially due to it having a small *T*_2_ value and thus less signal decay during STEAM’s TE^57^. Moreover, another study demonstrated improved detectability of both Lac and β-hydroxybutyrate using MEGA-sLASER compared to MEGA-PRESS^58^. Given that many of the commonly editable metabolites are present in lower concentrations in the human brain and can be challenging to detect at 3T, applying sLASER becomes valuable, offering enhanced reliability. It should be noted, however, that this increased reliability will depend on the metabolite(s) of interest.

The present results demonstrate that there should be an understanding that during study design, researchers should optimize the HERMES or HERCULES sequences, given that some metabolites may have poorer test-retest reliability. In other words, the sequences and parameters may need to be modified based on the metabolite(s) of interest in an experiment. Some studies have shown that multi-metabolite editing methods can be adapted as such. For example, Chan et al. (2016) devised the original HERMES approach specifically to target NAA and NAAG^24^, while Saleh et al. (2020) adapted HERMES to target ethanol, GABA, and GSH^59^. Likewise, another study modified HERMES to target Asp, NAA, and NAAG^60^. Other approaches include using multiplexed editing to detect GSH and Lac using either MEGA or DEW^61^. Furthermore, a distinct approach was employed in another study to investigate how PRESS and sLASER established biological relationships, finding strong agreement in both methods^62^. However, this may be dependent on the brain region. Altogether, the choice of multi-metabolite editing should be tailored to the metabolite(s) of interest pertinent to the study. This is particularly relevant for metabolite peaks that greatly overlap (e.g., NAA and NAAG) and whose subsequent individual quantification may be biased^62^.

The present study has several limitations. First, the order of the MRS acquisitions was not randomized or counterbalanced, potentially introducing order effects that could influence the study outcomes. Second, there was a difference in the water suppression schemes employed for the two localization methods. PRESS-localized scans utilized CHESS for water suppression, while sLASER scans used VAPOR, as recommended in recent consensus reports^30^. It is worth noting, though, that our implementation of PRESS lacked the capability to employ VAPOR. However, we chose to implement each localization with the recommended water suppression technique, aligning with standard practices in the community^32^. While reflecting real-world conditions, this decision introduces a potential source of variability between the two methods that should be considered when interpreting our results. Third, our TE of 82 ms was used in the present study, whereas the original HERMES and HERCULES papers used a TE of 80 ms^25,27^. Moreover, we were unable to assess the reliability of Asc. It will likely be worth examining the reliability of sLASER compared to PRESS in different brain or body regions as well. Specifically, there are loci of disease or treatment targets that may serve better for neurological and psychiatric disorders (e.g., the prefrontal cortex^11^). Another pertinent example is 2HG in brain tumors^63^.

In conclusion, this study has reported the test-retest reliability data for multi-metabolite editing using HERMES and HERCULES, in which volume localization was achieved with sLASER or PRESS. sLASER generally appears to be more reliable than PRESS in detecting and quantifying a variety of metabolites in a single acquisition. Yet, we recommend optimizing and adjusting multi-metabolite editing schemes to detect metabolites of interest. Given that there are a limited number of studies comparing PRESS and sLASER concomitantly, our research contributes to the efforts to support the reliability of MRS by examining two localization methods alongside two multi-metabolite editing techniques.

## Acknowledgments

This research was supported by the National Institute of Biomedical Imaging and Bioengineering of the National Institutes of Health under grant K99EB028828.

## Data Availability Statement

MRI and MRS data from this study are available on OpenNeuro (https://openneuro.org/datasets/ds005371). Note that the data of two participants was not uploaded because they did not provide the relevant consent. The data processing and statistical analysis code are available on GitHub (https://github.com/markmikkelsen/multi-metab-edit-reliability).

---

**Table S1.**
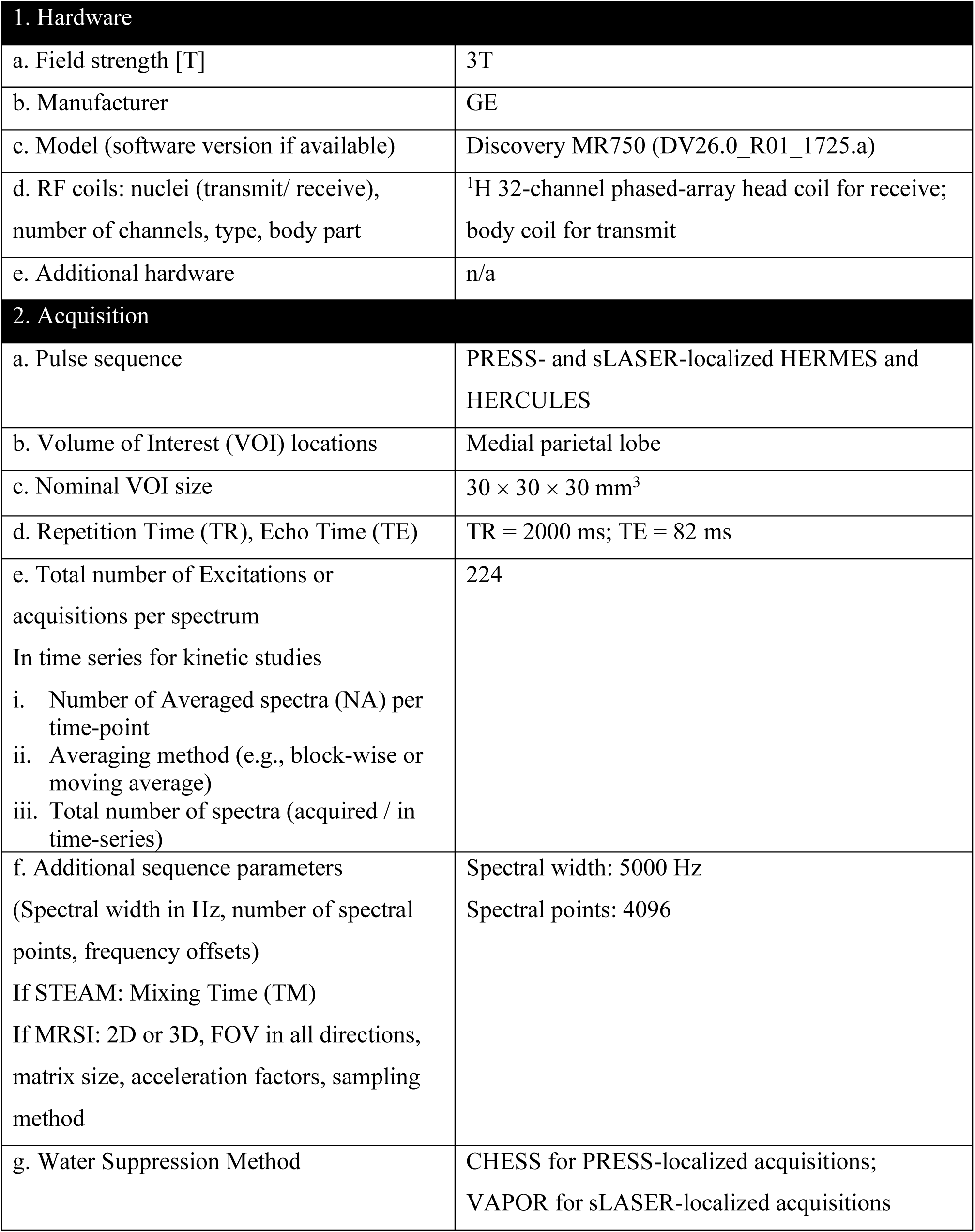
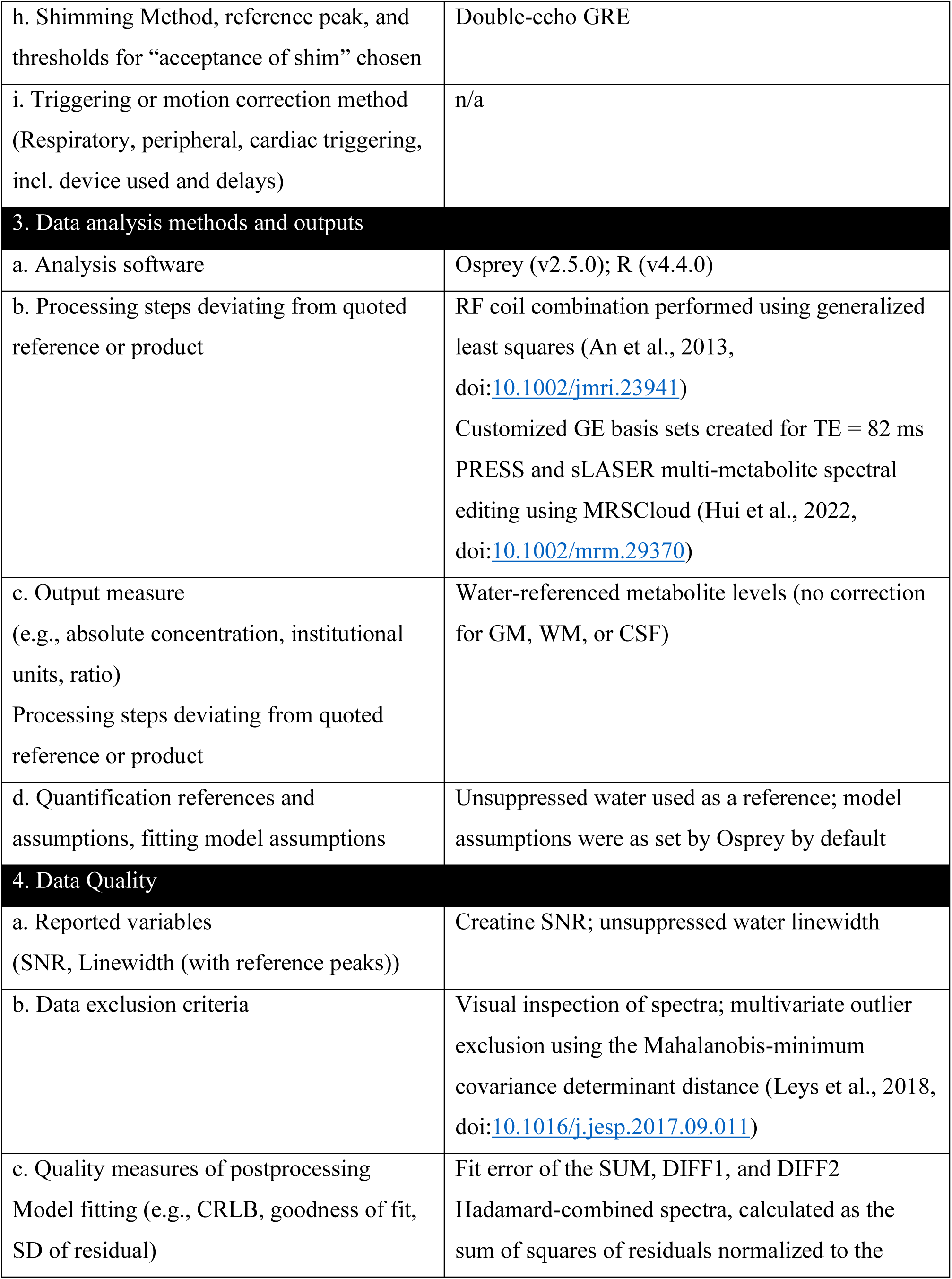
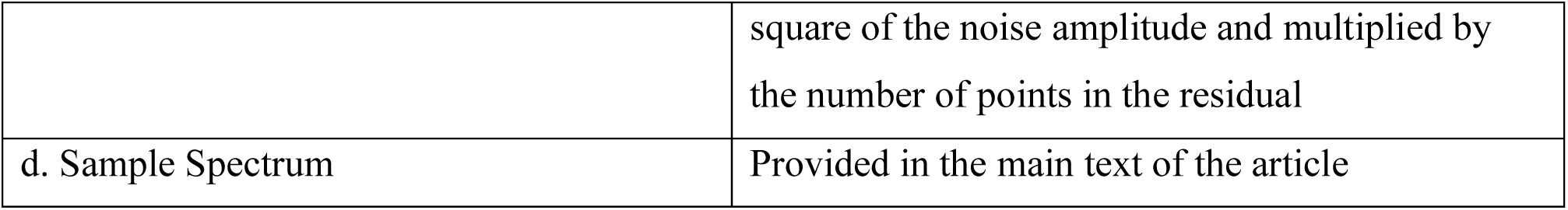
MRSinMRS checklist.

## References

1. Ip IB, Berrington A, Hess AT, Parker AJ, Emir UE, Bridge H. Combined fMRI-MRS acquires simultaneous glutamate and BOLD-fMRI signals in the human brain. Neuroimage. Jul 15 2017;155:113–119. doi:10.1016/j.neuroimage.2017.04.030

2. Yakovlev A, Gritskova A, Manzhurtsev A, et al. Dynamics of gamma-aminobutyric acid concentration in the human brain in response to short visual stimulation. MAGMA. Sep 16 2023;doi:10.1007/s10334-023-01118-7

3. DiNuzzo M, Mangia S, Moraschi M, Mascali D, Hagberg GE, Giove F. Perception is associated with the brain’s metabolic response to sensory stimulation. Elife. Feb 28 2022;11doi:10.7554/eLife.71016

4. Archibald J, MacMillan EL, Enzler A, Jutzeler CR, Schweinhardt P, Kramer JLK. Excitatory and inhibitory responses in the brain to experimental pain: A systematic review of MR spectroscopy studies. Neuroimage. Jul 15 2020;215:116794. doi:10.1016/j.neuroimage.2020.116794

5. Archibald J, MacMillan EL, Graf C, Kozlowski P, Laule C, Kramer JLK. Metabolite activity in the anterior cingulate cortex during a painful stimulus using functional MRS. Sci Rep. Nov 5 2020;10(1):19218. doi:10.1038/s41598-020-76263-3

6. Murray ME, Przybelski SA, Lesnick TG, et al. Early Alzheimer’s disease neuropathology detected by proton MR spectroscopy. J Neurosci. Dec 3 2014;34(49):16247–55. doi:10.1523/JNEUROSCI.2027-14.2014

7. Ford TC, Crewther DP. A Comprehensive Review of the (1)H-MRS Metabolite Spectrum in Autism Spectrum Disorder. Front Mol Neurosci. 2016;9:14. doi:10.3389/fnmol.2016.00014

8. Kim SY, Kaufman MJ, Cohen BM, et al. In Vivo Brain Glycine and Glutamate Concentrations in Patients With First-Episode Psychosis Measured by Echo Time-Averaged Proton Magnetic Resonance Spectroscopy at 4T. Biol Psychiatry. Mar 15 2018;83(6):484–491. doi:10.1016/j.biopsych.2017.08.022

9. Swanberg KM, Kurada AV, Prinsen H, Juchem C. Multiple sclerosis diagnosis and phenotype identification by multivariate classification of in vivo frontal cortex metabolite profiles. Sci Rep. Aug 16 2022;12(1):13888. doi:10.1038/s41598-022-17741-8

10. Chiu PW, Lui SSY, Hung KSY, et al. In vivo gamma-aminobutyric acid and glutamate levels in people with first-episode schizophrenia: A proton magnetic resonance spectroscopy study. Schizophr Res. Mar 2018;193:295–303. doi:10.1016/j.schres.2017.07.021

11. Ritter C, Buchmann A, Müller ST, et al. Evaluation of Prefrontal γ-Aminobutyric Acid and Glutamate Levels in Individuals With Major Depressive Disorder Using Proton Magnetic Resonance Spectroscopy. JAMA Psychiatry. Dec 1 2022;79(12):1209–1216. doi:10.1001/jamapsychiatry.2022.3384

12. Kara F, Joers JM, Deelchand DK, et al. (1)H MR spectroscopy biomarkers of neuronal and synaptic function are associated with tau deposition in cognitively unimpaired older adults. Neurobiol Aging. Apr 2022;112:16–26. doi:10.1016/j.neurobiolaging.2021.12.010

13. Graaf RAd. In Vivo NMR Spectroscopy. 3rd ed. Wiley; 2019.

14. Detcheverry F, Senthil S, Narayanan S, Badhwar A. Changes in levels of the antioxidant glutathione in brain and blood across the age span of healthy adults: A systematic review. Neuroimage Clin. 2023;40:103503. doi:10.1016/j.nicl.2023.103503

15. Porges EC, Jensen G, Foster B, Edden RA, Puts NA. The trajectory of cortical GABA across the lifespan, an individual participant data meta-analysis of edited MRS studies. Elife. Jun 1 2021;10doi:10.7554/eLife.62575

16. Bansal A, Simon MC. Glutathione metabolism in cancer progression and treatment resistance. J Cell Biol. Jul 2 2018;217(7):2291–2298. doi:10.1083/jcb.201804161

17. Kennedy L, Sandhu JK, Harper ME, Cuperlovic-Culf M. Role of Glutathione in Cancer: From Mechanisms to Therapies. Biomolecules. Oct 9 2020;10(10)doi:10.3390/biom10101429

18. Li H, Heise KF, Chalavi S, Puts NAJ, Edden RAE, Swinnen SP. The role of MRS-assessed GABA in human behavioral performance. Prog Neurobiol. May 2022;212:102247. doi:10.1016/j.pneurobio.2022.102247

19. Schur RR, Draisma LW, Wijnen JP, et al. Brain GABA levels across psychiatric disorders: A systematic literature review and meta-analysis of (1) H-MRS studies. Hum Brain Mapp. Sep 2016;37(9):3337–52. doi:10.1002/hbm.23244

20. Thomson AR, Pasanta D, Arichi T, Puts NA. Neurometabolite differences in Autism as assessed with Magnetic Resonance Spectroscopy: A systematic review and meta-analysis. Neurosci Biobehav Rev. Jul 2024;162:105728. doi:10.1016/j.neubiorev.2024.105728

21. Quadrelli S, Mountford C, Ramadan S. Systematic review of in-vivo neuro magnetic resonance spectroscopy for the assessment of posttraumatic stress disorder. Psychiatry Res Neuroimaging. Dec 30 2018;282:110–125. doi:10.1016/j.pscychresns.2018.07.001

22. McKiernan E, Su L, O’Brien J. MRS in neurodegenerative dementias, prodromal syndromes and at-risk states: A systematic review of the literature. NMR Biomed. Jul 2023;36(7):e4896. doi:10.1002/nbm.4896

23. Sarlo GL, Holton KF. Brain concentrations of glutamate and GABA in human epilepsy: A review. Seizure. Oct 2021;91:213–227. doi:10.1016/j.seizure.2021.06.028

24. Chan KL, Puts NA, Schar M, Barker PB, Edden RA. HERMES: Hadamard encoding and reconstruction of MEGA-edited spectroscopy. Magn Reson Med. Jul 2016;76(1):11–9. doi:10.1002/mrm.26233

25. Saleh MG, Oeltzschner G, Chan KL, et al. Simultaneous edited MRS of GABA and glutathione. Neuroimage. Nov 15 2016;142:576–582. doi:10.1016/j.neuroimage.2016.07.056

26. Chan KL, Oeltzschner G, Saleh MG, Edden RAE, Barker PB. Simultaneous editing of GABA and GSH with Hadamard-encoded MR spectroscopic imaging. Magn Reson Med. Jul 2019;82(1):21–32. doi:10.1002/mrm.27702

27. Oeltzschner G, Saleh MG, Rimbault D, et al. Advanced Hadamard-encoded editing of seven low-concentration brain metabolites: Principles of HERCULES. Neuroimage. Jan 15 2019;185:181–190. doi:10.1016/j.neuroimage.2018.10.002

28. Bottomley PA. Spatial localization in NMR spectroscopy in vivo. Ann N Y Acad Sci. 1987;508:333–48. doi:10.1111/j.1749-6632.1987.tb32915.x

29. Scheenen TWJ, Klomp DWJ, Wijnen JP, Heerschap A. Short echo time 1H-MRSI of the human brain at 3T with minimal chemical shift displacement errors using adiabatic refocusing pulses. Magn Reson Med. 2008;59(1):1–6. doi:10.1002/mrm.21302

30. Wilson M, Andronesi O, Barker PB, et al. Methodological consensus on clinical proton MRS of the brain: Review and recommendations. Magn Reson Med. Aug 2019;82(2):527–550. doi:10.1002/mrm.27742

31. Oz G, Deelchand DK, Wijnen JP, et al. Advanced single voxel (1) H magnetic resonance spectroscopy techniques in humans: Experts’ consensus recommendations. NMR Biomed. Jan 10 2020:e4236. doi:10.1002/nbm.4236

32. Tkac I, Deelchand D, Dreher W, et al. Water and lipid suppression techniques for advanced 1H MRS and MRSI of the human brain: Experts’ consensus recommendations. NMR Biomed. May 2021;34(5):e4459. doi:10.1002/nbm.4459

33. Lin A, Andronesi O, Bogner W, et al. Minimum Reporting Standards for in vivo Magnetic Resonance Spectroscopy (MRSinMRS): Experts’ consensus recommendations. NMR Biomed. May 2021;34(5):e4484. doi:10.1002/nbm.4484

34. Oeltzschner G, Zollner HJ, Hui SCN, et al. Osprey: Open-source processing, reconstruction & estimation of magnetic resonance spectroscopy data. J Neurosci Methods. Sep 1 2020;343:108827. doi:10.1016/j.jneumeth.2020.108827

35. An L, Willem van der Veen J, Li S, Thomasson DM, Shen J. Combination of multichannel single-voxel MRS signals using generalized least squares. J Magn Reson Imaging. Jun 2013;37(6):1445–50. doi:10.1002/jmri.23941

36. Klose U. In vivo proton spectroscopy in presence of eddy currents. Magn Reson Med. Apr 1990;14(1):26–30. doi:10.1002/mrm.1910140104

37. Mikkelsen M, Tapper S, Near J, Mostofsky SH, Puts NAJ, Edden RAE. Correcting frequency and phase offsets in MRS data using robust spectral registration. NMR Biomed. Oct 2020;33(10):e4368. doi:10.1002/nbm.4368

38. Barkhuijsen H, de Beer R, van Ormondt D. Improved algorithm for noniterative time-domain model fitting to exponentially damped magnetic resonance signals. Journal of Magnetic Resonance (1969). 1987/07/01/ 1987;73(3):553–557. doi:10.1016/0022-2364(87)90023-0

39. Hui SCN, Saleh MG, Zollner HJ, et al. MRSCloud: A cloud-based MRS tool for basis set simulation. Magn Reson Med. Nov 2022;88(5):1994–2004. doi:10.1002/mrm.29370

40. Zhang Y, An L, Shen J. Fast computation of full density matrix of multispin systems for spatially localized in vivo magnetic resonance spectroscopy. Med Phys. Aug 2017;44(8):4169–4178. doi:10.1002/mp.12375

41. Landheer K, Swanberg KM, Juchem C. Magnetic resonance Spectrum simulator (MARSS), a novel software package for fast and computationally efficient basis set simulation. NMR Biomed. May 2021;34(5):e4129. doi:10.1002/nbm.4129

42. Leys C, Klein O, Dominicy Y, Ley C. Detecting multivariate outliers: Use a robust variant of the Mahalanobis distance. Journal of Experimental Social Psychology. 2018/01/01/ 2018;74:150–156. doi:10.1016/j.jesp.2017.09.011

43. Nacewicz BM, Angelos L, Dalton KM, et al. Reliable non-invasive measurement of human neurochemistry using proton spectroscopy with an anatomically defined amygdala-specific voxel. Neuroimage. Feb 1 2012;59(3):2548–59. doi:10.1016/j.neuroimage.2011.08.090

44. Baeshen A, Wyss PO, Henning A, et al. Test-Retest Reliability of the Brain Metabolites GABA and Glx With JPRESS, PRESS, and MEGA-PRESS MRS Sequences in vivo at 3T. J Magn Reson Imaging. Apr 2020;51(4):1181–1191. doi:10.1002/jmri.26921

45. Lally N, An L, Banerjee D, et al. Reliability of 7T (1) H-MRS measured human prefrontal cortex glutamate, glutamine, and glutathione signals using an adapted echo time optimized PRESS sequence: A between- and within-sessions investigation. J Magn Reson Imaging. Jan 2016;43(1):88–98. doi:10.1002/jmri.24970

46. Liu XL, Li L, Li JN, Rong JH, Liu B, Hu ZX. Reliability of Glutamate Quantification in Human Nucleus Accumbens Using Proton Magnetic Resonance Spectroscopy at a 70-cm Wide-Bore Clinical 3T MRI System. Front Neurosci. 2017;11:686. doi:10.3389/fnins.2017.00686

47. Napolitano A, Kockenberger W, Auer DP. Reliable gamma aminobutyric acid measurement using optimized PRESS at 3 T. Magn Reson Med. Jun 2013;69(6):1528–33. doi:10.1002/mrm.24397

48. Fayed N, Modrego PJ, Medrano J. Comparative test-retest reliability of metabolite values assessed with magnetic resonance spectroscopy of the brain. The LCModel versus the manufacturer software. Neurol Res. Jun 2009;31(5):472–7. doi:10.1179/174313209x395481

49. Shyu C, Elsaid S, Truong P, Chavez S, Le Foll B. MR Spectroscopy of the Insula: Within- and between-Session Reproducibility of MEGA-PRESS Measurements of GABA+ and Other Metabolites. Brain Sci. Nov 19 2021;11(11)doi:10.3390/brainsci11111538

50. Duda JM, Moser AD, Zuo CS, et al. Repeatability and reliability of GABA measurements with magnetic resonance spectroscopy in healthy young adults. Magn Reson Med. May 2021;85(5):2359–2369. doi:10.1002/mrm.28587

51. Hupfeld KE, Zöllner HJ, Hui SCN, et al. Impact of acquisition and modeling parameters on the test-retest reproducibility of edited GABA. NMR Biomed. Apr 2024;37(4):e5076. doi:10.1002/nbm.5076

52. Prisciandaro JJ, Mikkelsen M, Saleh MG, Edden RAE. An evaluation of the reproducibility of (1)H-MRS GABA and GSH levels acquired in healthy volunteers with J-difference editing sequences at varying echo times. Magn Reson Imaging. Jan 2020;65:109–113. doi:10.1016/j.mri.2019.10.004

53. Elsaid S, Truong P, Sailasuta N, Le Foll B. Evaluating Back-to-Back and Day-to-Day Reproducibility of Cortical GABA+ Measurements Using Proton Magnetic Resonance Spectroscopy ((1)H MRS). Int J Mol Sci. Apr 23 2023;24(9)doi:10.3390/ijms24097713

54. Brix MK, Ersland L, Hugdahl K, et al. Within- and between-session reproducibility of GABA measurements with MR spectroscopy. J Magn Reson Imaging. Aug 2017;46(2):421–430. doi:10.1002/jmri.25588

55. Deelchand DK, Kantarci K, Oz G. Improved localization, spectral quality, and repeatability with advanced MRS methodology in the clinical setting. Magn Reson Med. Mar 2018;79(3):1241–1250. doi:10.1002/mrm.26788

56. Marsman A, Boer VO, Luijten PR, Hulshoff Pol HE, Klomp DWJ, Mandl RCW. Detection of Glutamate Alterations in the Human Brain Using (1)H-MRS: Comparison of STEAM and sLASER at 7 T. Front Psychiatry. 2017;8:60. doi:10.3389/fpsyt.2017.00060

57. Okada T, Kuribayashi H, Kaiser LG, et al. Repeatability of proton magnetic resonance spectroscopy of the brain at 7 T: effect of scan time on semi-localized by adiabatic selective refocusing and short-echo time stimulated echo acquisition mode scans and their comparison. Quantitative Imaging in Medicine and Surgery. 2020;11(1):9–20.

58. Dacko M, Lange T. Improved detection of lactate and beta-hydroxybutyrate using MEGA-sLASER at 3 T. NMR Biomed. Jul 2019;32(7):e4100. doi:10.1002/nbm.4100

59. Saleh MG, Wang M, Mikkelsen M, et al. Simultaneous edited MRS of GABA, glutathione, and ethanol. NMR Biomed. Apr 2020;33(4):e4227. doi:10.1002/nbm.4227

60. Chan KL, Saleh MG, Oeltzschner G, Barker PB, Edden RAE. Simultaneous measurement of Aspartate, NAA, and NAAG using HERMES spectral editing at 3 Tesla. Neuroimage. Jul 15 2017;155:587–593. doi:10.1016/j.neuroimage.2017.04.043

61. Chan KL, Snoussi K, Edden RAE, Barker PB. Simultaneous detection of glutathione and lactate using spectral editing at 3 T. NMR Biomed. Dec 2017;30(12)doi:10.1002/nbm.3800

62. Hui SCN, Zöllner HJ, Gong T, et al. sLASER and PRESS perform similarly at revealing metabolite-age correlations at 3 T. Magn Reson Med. Feb 2024;91(2):431–442. doi:10.1002/mrm.29895

63. Berrington A, Voets NL, Plaha P, et al. Improved localisation for 2-hydroxyglutarate detection at 3T using long-TE semi-LASER. Tomography. Jun 2016;2(2):94–105. doi:10.18383/j.tom.2016.00139

